# Is there a placental microbiota? A critical review and re-analysis of published placental microbiota datasets

**DOI:** 10.1101/2022.07.18.500562

**Authors:** Jonathan Panzer, Roberto Romero, Jonathan M. Greenberg, Andrew D. Winters, Jose Galaz, Nardhy Gomez-Lopez, Kevin R. Theis

## Abstract

The existence of a placental microbiota is under debate. The human placenta has historically been considered sterile and microbial colonization has been associated with adverse pregnancy outcomes. Yet, recent investigations using DNA sequencing reported a microbiota in human placentas from typical term pregnancies. However, this detected microbiota could represent background DNA contamination. Using fifteen publicly available 16S rRNA gene datasets, existing data were uniformly re-analyzed. 16S rRNA gene Amplicon Sequence Variants (ASVs) identified as *Lactobacillus* were highly abundant in eight of fifteen studies. However, the prevalence of *Lactobacillus*, a typical vaginal bacterium, was clearly driven by bacterial contamination from vaginal delivery and background DNA. After removal of likely DNA contaminants, *Lactobacillus* ASVs were highly abundant in only one of five studies for which data analysis could be restricted to placentas from term cesarean deliveries. A six study sub-analysis targeting the 16S rRNA gene V4 hypervariable region demonstrated that bacterial profiles of placental samples and technical controls share principal bacterial ASVs and that placental samples clustered primarily by study origin and mode of delivery. Across studies, placentas from typical term pregnancies did not share a consistent bacterial taxonomic signal. Contemporary DNA-based evidence does not support the existence of a placental microbiota.

**IMPORTANCE:** Early-gestational microbial influences on human development are unclear. By applying DNA sequencing technologies to placental tissue, bacterial DNA signals were observed, leading some to conclude that a live bacterial placental microbiome exists in typical term pregnancy. However, the low-biomass nature of the proposed microbiome and high sensitivity of current DNA sequencing technologies indicate that the signal may alternatively derive from environmental or delivery-associated bacterial DNA contamination. Here we address these alternatives with a re- analysis of 16S rRNA gene sequencing data from 15 publicly available placental datasets. After identical DADA2 pipeline processing of the raw data, subanalyses were performed to control for mode of delivery and environmental DNA contamination. Both environment and mode of delivery profoundly influenced the bacterial DNA signal from term-delivered placentas. Aside from these contamination-associated signals, consistency was lacking across studies. Thus, placentas delivered at term are unlikely to be the original source of observed bacterial DNA signals.

## INTRODUCTION

The womb has historically been considered sterile throughout typical pregnancy (1–3); yet, the detection of microorganisms, especially bacteria, in some placentas from complicated pregnancies is an established phenomenon (4–7). For instance, there are demonstrated associations between bacterial colonization of the placenta and preterm labor (5, 8–15), preterm prelabor rupture of membranes (PPROM) (11, 12), histological chorioamnionitis (8, 14, 16), and clinical chorioamnionitis (12–14, 16). Therefore, research has largely focused on the presence (9, 17–20) and types of bacteria (21–26) associated with the human placenta in the context of spontaneous preterm birth and other pregnancy complications.

However, in 2014, bacterial DNA-based evidence was presented for a universal low-biomass placental microbiota even among placentas from term pregnancies (27). Since placental colonization by bacteria suggests that fetal colonization is also feasible, this study revitalized the *in utero* colonization hypothesis, which maintains that commensal bacteria residing in the placenta and/or amniotic fluid colonize the developing fetus during gestation (3, 28–34). The *in utero* colonization hypothesis stands in stark contrast to the traditional sterile womb hypothesis, which posits fetal sterility until colonization at delivery or following rupture of the amniotic membranes (1-3, 35-41). Since publication of this seminal study in 2014, many other studies have similarly utilized DNA sequencing techniques to investigate the existence of a placental microbiota in term pregnancies (28, 29, 34, 36–65). Yet, the existence of a placental microbiota remains under debate eight years later (66–69).

The debate over the existence of a placental microbiota is fueled by several issues. First, the placenta cannot be readily sampled *in utero*. Thus, attempts at characterizing a placental microbiota have entailed collection of placental samples following either a vaginal or cesarean delivery. While both delivery methods can introduce bacterial contamination (36, 38, 40, 42, 51, 70), in the form of vaginal and skin bacteria, respectively, vaginal delivery likely exposes the placenta to bacterial contamination to an extent that would overwhelm any weak bacterial DNA signal legitimately present in placental tissue *in utero* (40, 42, 56). Thus, to establish that a placental microbiota exists, it must be documented in placentas from term cesarean deliveries to minimize misinterpretation of potential delivery associated contamination (3, 29, 71).

Second, a lack of robust technical controls has made it difficult to determine if reagent or environmental DNA contamination might be the source of bacterial DNA signals attributed to placentas rather than a resident placental microbiota (27, 29, 46, 53, 54, 57–61, 63–65), given that such a theoretically sparse bacterial community could easily be obfuscated by background DNA contamination in laboratories, kits, and reagents (39, 72–75). Technical controls and sterile collection conditions are therefore essential for the verification of a placental microbiota. Indeed, several recent studies have shown that the bacterial loads (41) and profiles of placentas from term cesarean deliveries do not exceed or differ from those of technical controls (41, 42).

Finally, a lack of consistency in the analytical pipelines used to process the DNA sequence data has resulted in additional debate, including how sequences should be grouped or split into taxonomic units (73, 76, 77). Specifically, too coarse or too fine a taxonomic resolution could either potentially reveal a shared bacterial DNA signal between placental tissues and technical controls or a signal unique to the placenta, respectively.

Ultimately, if there is a placental microbiota it should exist in a majority of, if not all, placentas from women delivering at term without complications, and there should be some degree of consistency in the bacterial taxa residing in placentas across studies. For example, the human vaginal microbiota worldwide is consistently predominated by various species of *Lactobacillus* and, in a smaller proportion of women, higher diversity bacterial communities exist, which consist of nevertheless predictable genera such as *Prevotella*, *Sneathia*, *Megasphaera*, *Atopobium*, *Mobiluncus*, *Streptococcus*, and *Gardnerella* (78, 79). Yet, among investigators proposing the existence of a placental microbiota, there are conflicting reports regarding its predominant bacterial members (27-29, 44, 49, 54, 58, 62-65) and, when complementary culture results are available, placental samples are often culture negative or the bacteria recovered are discrepant with the DNA sequencing results (19, 28, 40, 41, 44, 56, 80–85).

Given these current conflicting conclusions regarding the existence of a placental microbiota, here we performed a critical review and re-analysis of fifteen publicly available 16S rRNA gene sequencing datasets from human placental microbiota studies for which sample distinguishing metadata were available (29, 36–44, 50, 53, 57, 86). In this re-analysis we standardized the analytical process to enable assessment of taxonomic consistency in placental bacterial profiles across studies conducted by different laboratories across the world **(Figure S1)**. Briefly, raw sequencing data from each study were processed using the same analytical pipeline, the Divisive Amplicon Denoising Algorithm (DADA2), to provide consistent sequence filtering and clustering methods (87). Additionally, eight of the fifteen studies included sequence data for at least six technical controls to account for potential background DNA contamination (88). For these studies, the R package DECONTAM (88) was used to identify and remove likely DNA contaminants and we report the DNA signal from the bacterial taxa that remained.

Three primary analyses were performed. The first analysis was a comparison of the bacterial profiles of placental samples to technical controls for studies which included at least six controls for background DNA contamination (88) since this environmental contamination could be a source of the purported placental bacterial DNA signal. Ideally, a valid placental microbiota would be expected to exhibit a bacterial DNA signal distinct from that of kit reagents or surrounding laboratory environments. The first analysis revealed no consistent differentiation between the bacterial DNA signal from placental samples and technical controls, or vaginal swabs when available.

The second analysis was restricted to placentas from term cesarean deliveries so as to avoid potential bacterial contamination of placentas that could occur during vaginal delivery (36, 42, 51, 89). If there were a placental microbiota, the bacterial DNA signals should be clear and consistent across placentas from term cesarean deliveries. This analysis was therefore performed using data from the six studies for which placental samples could be restricted to those from term cesarean deliveries. Of the studies which included at least six background technical controls, after likely contaminant removal, there was no consistent bacterial DNA signal among placental samples.

The third and final analysis was restricted to studies that targeted the V4 hypervariable region of the 16S rRNA gene to control for any variation which might arise due to variation in targeted 16S rRNA gene hypervariable regions across studies or the DNA sequence processing algorithms used. A valid placental microbiota would be expected to be independent of study identity and mode of delivery; however, both of these factors largely contributed to the structure of placental bacterial profiles. Indeed, in these studies, there was a large degree of similarity in the bacterial profiles of placental samples and technical controls from the same study, and there were no bacterial taxa that were consistently identified across studies whose presence could not be explained by background DNA contamination.

Collectively, these analyses do not support the presence of a placental microbiota in typical term pregnancies. Observed bacterial signals were products of mode of delivery and background DNA contamination. Although there may be a true, consistent, extremely low biomass bacterial signal beyond the limits of detection by contemporary 16S rRNA gene sequencing, it remains to be demonstrated that the placenta harbors a legitimate bacterial DNA signal or a viable microbiota in typical human pregnancy.

## RESULTS

### Overview of studies included in this re-analysis

Fifteen studies **(Table 1)** were included in this re-analysis of investigations of the existence of a placental microbiota. Seven included sequence data from the V4 hypervariable region of the 16S rRNA gene (29, 39, 41, 43, 44, 50, 86), allowing for direct comparisons of sequence data across six of those studies (29, 39, 41, 44, 50, 86); one study could not be included in the direct comparison due to short read lengths of sequences in the publicly available dataset (43). Three of the remaining studies included sequence data from the V1-V2 16S rRNA gene hypervariable region (36–38), two studies sequenced the V6-V8 region (40, 53), and one study each sequenced the V3-V4 (58), V4-V5 (57), and V5-V7 (42) regions. All fifteen studies included at least one placental sample from a term cesarean delivery, but only eight included more than one term cesarean delivered placenta and sufficient background technical controls [i.e., N = 6 (88)] to infer likely DNA contaminants using the R package DECONTAM (36, 39–43, 50, 86) (**Figure 2**). Two of these studies lacked available metadata to discriminate placental samples by gestational age at delivery (42, 50), leaving a total of six studies (36, 39–41, 43, 86) for assessing uniformity of bacterial profiles among term cesarean delivered placentas across studies while accounting for potential background DNA contamination (**Figure 2**). Notably, five of these six studies concluded that there was no evidence for a placental microbiota in uncomplicated term pregnancies (36, 39–41, 86) (**Figure 2**). In contrast, the four studies which did not include sequence data from background technical controls concluded that a placental microbiota does exist (29, 53, 57, 58) (**Figure 2**).

**Figure 1.**
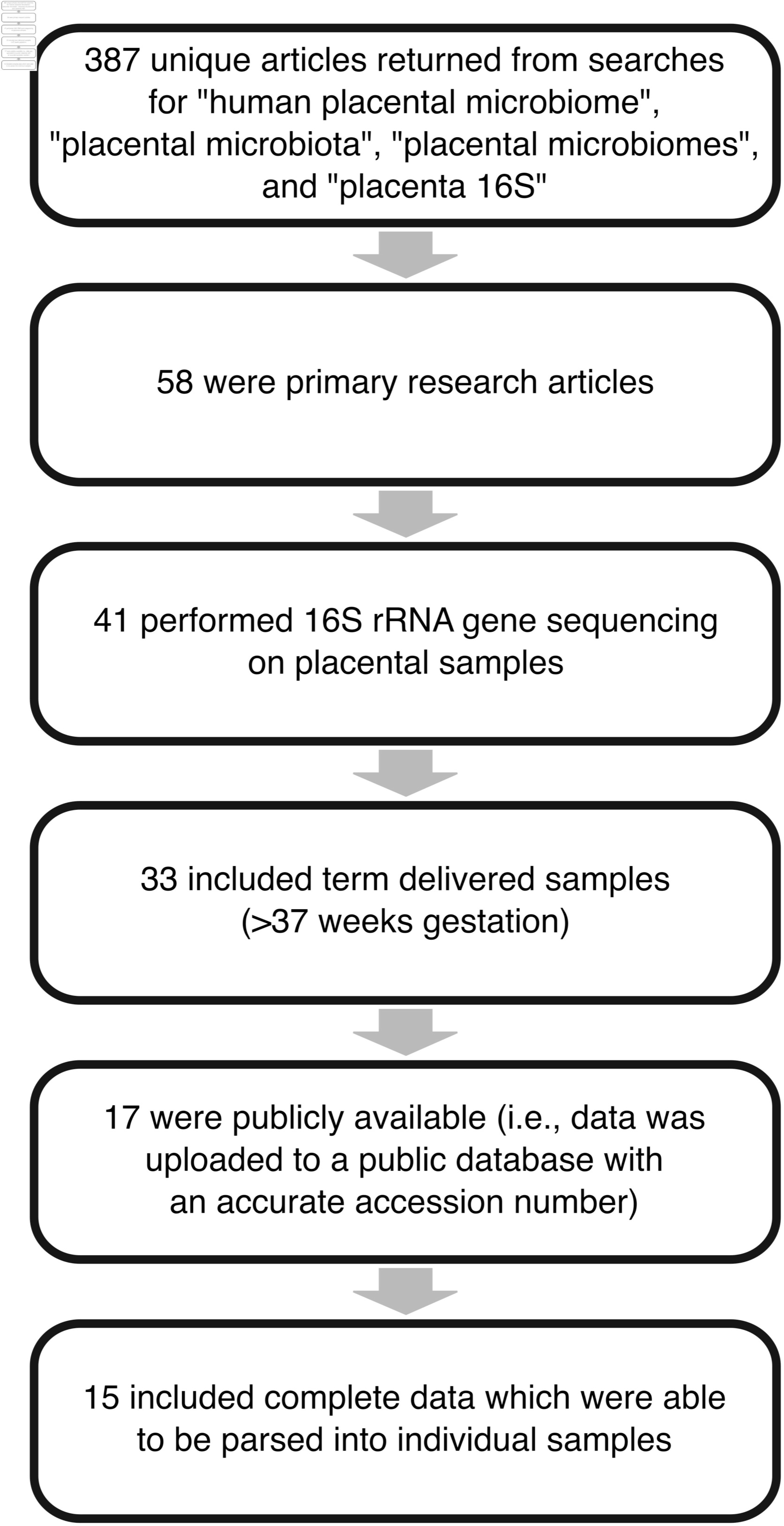
Study inclusion flowchart Four searches were performed on PubMed to identify studies for inclusion in the re-analysis. Filtering criteria were: primary research article, 16S rRNA gene sequencing data, placentas obtained from term deliveries, sequencing data accessible with published accession number, and sufficient metadata available to parse sequencing data into individual samples.

**Figure 2.**
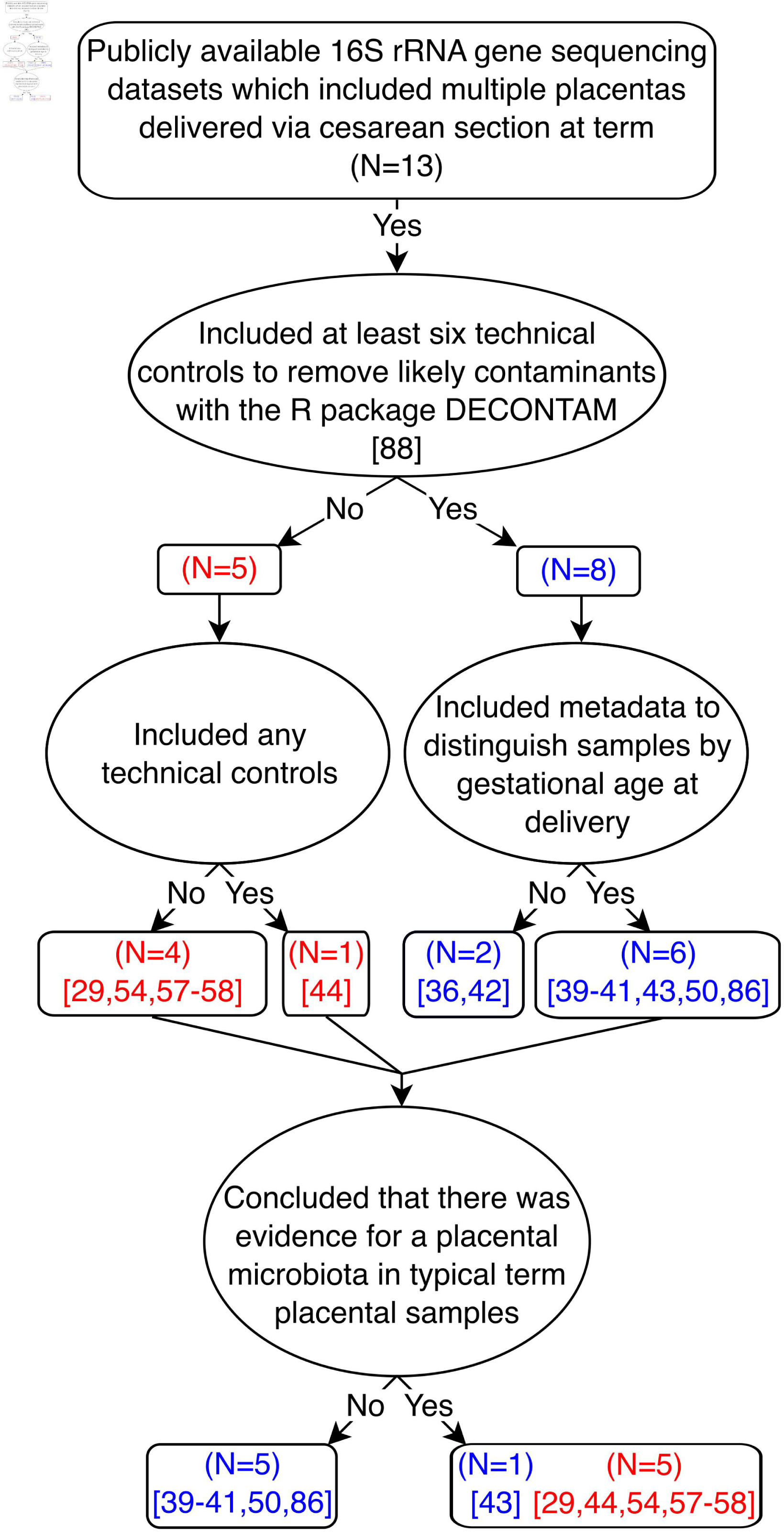
Conclusions of thirteen studies evaluating the existence of a placental microbiota, which included data from multiple placentas delivered via cesarean section at term. The studies are principally separated and contrasted depending upon whether they included technical controls to account for potential background DNA contamination.

**Table 1.** Overview of placental microbiota studies that were based on 16S rRNA gene sequencing data and that were included in this critical review and re-analysis. The presented study characteristics include the: name of the first author(s); geographical location at which subjects were sampled; specific 16S rRNA gene hypervariable region that was targeted for sequencing; number of placentas sampled by mode of delivery and whether delivery was before (i.e., preterm) or after (i.e. term) 37 weeks; number of technical controls included to address potential background DNA contamination; and whether we were able to categorize placental samples based on mode of delivery, gestational age at delivery (i.e., before or after 37 weeks), and whether DECONTAM analysis could be performed to identify background DNA contaminants (i.e., N ≥ 6 technical controls included in the study) (88). Square brackets indicate that available sample metadata did not allow for placentas to be grouped by gestational age at delivery.

| Study | Geographic Location | 16S rRNA gene hypervariable region | Cesarean |  | Vaginal |  | Technical Controls | Ability to group by |  | Ability to run DECONTAM |
| --- | --- | --- | --- | --- | --- | --- | --- | --- | --- | --- |
|  |  |  | Term | Preterm | Term | Preterm |  | Delivery | Gestational Age |  |
| de Goffau Part I<sup>b</sup> | Cambridge, UK | V1-V2 | 80 | 0 | 0 | 0 | 47 | X | X | X |
| Lauder | Philadelphia, PA, USA | V1-V2 | 1 | 0 | 5 | 0 | 39 | X | X | X |
| Leiby | Philadelphia, PA, USA | V1-V2 | 1 | 5 | 19 | 15 | 103 | X | X | X |
| Olomu | Lansing, MI, USA | V4 | 47<sup>a</sup> | 0 | 0 | 0 | 131 | X | X | X |
| Parnell | St. Louis, MO, USA | V4 | 34 | 0 | 23 | 0 | 21 | X | X | X |
| Theis<sup>c</sup> | Detroit, MI, USA | V4 | 29<sup>a</sup> | 0 | 0 | 0 | 43 | X | X | X |
| Theis, Winters | Detroit, MI, USA | V4 | 28 | 14 | 21 | 6 | 12 | X | X | X |
| Sterpu | Stockholm, Sweden | V6-V8 | 50<sup>a</sup> | 0 | 26 | 0 | 6 | X | X | X |
| Dinsdale | South Hedland, Australia | V4 | [19] |  | [31] |  | 8 | X |  | X |
| Tang | Shanghai, China | V3-V4 | 15<sup>a</sup> | 0 | 0 | 0 | 0 | X | X |  |
| Liu | Kunming, China | V4-V5 | 42 | 0 | 36 | 0 | 0 | X | X |  |
| Gomez-Arango | Brisbane, Australia | V6-V8 | 16<sup>a</sup> | 0 | 20 | 0 | 0 | X | X |  |
| Younge | Durham, NC, USA | V4 | 5<sup>a</sup> | 5 | 0 | 0 | 0 | X |  |  |
| Leon | London, England, UK | V5-V7 | [136] |  | [120] |  | 21 |  |  | X |
| Seferovic | Houston, TX, USA | V4 | 26 | 8 | 0 | 18 | 2 |  |  |  |

<sup>a</sup> Indicates that placental samples were delivered without labor.
<sup>b</sup> Analyzed data are from the Cohort 1 component of the study.
<sup>c</sup> Analyzed data are from the nested PCR component of the study.

### *Lactobacillus* ASVs are the most consistently identified ASVs across placental microbiota studies

After processing the raw 16S rRNA gene sequence data from placental samples from all 15 studies through the same DADA2 analytical pipeline, the most prominent bacterial ASVs, as defined by mean relative abundance, across studies were classified as *Lactobacillus*, *Escherichia/Shigella*, *Staphylococcus*, *Streptococcus*, and *Pseudomonas* **(Table S1)**.

*Lactobacillus* ASVs were among the top five ranked ASVs in eight of the 15 studies (37-39, 42, 50, 53, 57, 86), making *Lactobacillus* the most consistently detected genus in placental samples across studies with publicly available 16S rRNA gene sequencing data.

The detection of *Lactobacillus* ASVs was not exclusive to the targeted sequencing of specifically any one 16S rRNA gene hypervariable region; *Lactobacillus* ASVs were found among the top five ASVs in the dataset of at least one study targeting the V1-V2, V4, V4-V5, V5-V7, or V6-V8 hypervariable region(s) of the 16S rRNA gene **(Table S1)**. Other genera which were not 16S rRNA gene hypervariable region specific and were detected in the top five ranked ASVs in more than one dataset, but in no more than four, included *Staphylococcus* (40, 44, 54, 86), *Streptococcus* (38, 40, 42), and *Pseudomonas* (50, 54, 57). In contrast, *Escherichia/Shigella* ASVs were exclusively among the top five ranked ASVs in datasets of studies that targeted the V4 hypervariable region of the 16S rRNA gene for sequencing (3/7 such datasets) (29, 39, 86) **(Table S1)**.

### *Lactobacillus* ASVs in placental samples are typically contaminants introduced through vaginal delivery and/or background DNA contamination

While it can be difficult to identify the definitive source of a particular ASV in placental samples, the difference in *Lactobacillus* predominance between vaginally delivered placentas and cesarean delivered placentas is striking. *Lactobacillus* ASVs were among the top five ASVs in five of seven datasets which included placentas from vaginal deliveries before running the R package DECONTAM, and three of four datasets which included placentas from vaginal deliveries after running DECONTAM **(Table 2, Table S1)**. Consider, for instance, the Lauder et al. (37) and Leiby et al. (38) datasets. While all samples in the Lauder et al. dataset (37) had *Lactobacillus* ASVs, the percentage of *Lactobacillus* normalized reads in cesarean delivered placental samples was 23% compared to 46% in vaginally delivered placental samples. In the Leiby et al. dataset (38) only four of 23 (17%) cesarean delivered placentas had any *Lactobacillus* ASVs, and they made up only 2% of the total reads from their respective samples. In contrast, 35 of 116 (30%) placentas from vaginal deliveries had *Lactobacillus* ASVs, and they made up 22% of the total reads from those 35 samples.

**Table 2.** Summary of prominent bacterial ASVs in term cesarean delivered placental samples before and after removal of background DNA contaminants using the R package DECONTAM. The top five ASVs as determined by mean relative abundance across placental samples after DADA2 processing are provided for studies which could be restricted to cesarean delivered placental samples. Asterisks indicate ASV sequence genus level classifications which were assigned by NCBI BLAST with the highest percent identity in excess of 95%. The Liu et al. [1] dataset could not be assessed post-DECONTAM since no technical controls were included in the study.

| Study | 16S rRNA<br>gene<br>hypervariable<br>region | ASV1 | ASV2 | ASV3 | ASV4 | ASV5 |
| --- | --- | --- | --- | --- | --- | --- |
|  |  | **Before DECONTAM** |  |  |  |  |
| Olomu | V4 | *Escherichia/Shigella* | ***Lactobacillus*** | ***Lactobacillus*** | *Finegoldia* | *Fenollaria* |
| Parnell | V4 | *Leptospira* | *Leptospira* | *Leptospira* | *Cutibacterium* | *Leptospira* |
| Theis | V4 | *Achromobacter* | *Delftia* | *Phyllobacterium* | *Clostridium sensu stricto* 5 | *Stenotrophomonas* |
| Theis, Winters | V4 | ***Lactobacillus*** | *Staphylococcus* | *Escherichia/Shigella* | *Serratia* * | *Streptococcus* |
| Liu | V4-V5 | *Pseudomonas* | *Pantoea* * | *Pantoea* * | ***Lactobacillus*** | *Pantoea* * |
| Sterpu | V6-V8 | *Cutibacterium* | *Staphylococcus* | *Streptococcus* | *Streptococcus* | *Streptococcus* |
|  |  | **After DECONTAM** |  |  |  |  |
| Olomu | V4 | *Fenollaria* | *Acinetobacter* | ***Lactobacillus*** | *Campylobacter* | *Peptoniphilus* |
| Parnell | V4 | *Leptospira* | *Leptospira* | *Leptospira* | *Cutibacterium* | *Leptospira* |
| Theis | V4 | *Achromobacter* | Alcaligenaceae | *Escherichia/Shigella* | *Achromobacter* | Alcaligenaceae |
| Theis, Winters | V4 | *Corynebacterium* | *Serratia* * | *Mycoplasma* | *Corynebacterium* | *Fusobacterium* |
| Sterpu | V6-V8 | *Staphylococcus* | *Gardnerella* | *Staphylococcus* | *Staphylococcus* | *Streptococcus* |

ASV1-5 are rank designations based on percent relative abundance

*Lactobacillus* ASVs were among the top five ranked ASVs in three (39, 57, 86) of the six datasets which could be restricted to placental samples from cesarean term deliveries **(Table 2)**. Yet, after removing potential background DNA contaminants using DECONTAM, only the Olomu et al. (39) dataset still retained a *Lactobacillus* ASV in the top five ranked ASVs **(Table 2)**. Notably, the authors of that study identified the origin of *Lactobacillus* in placental samples as well-to-well DNA contamination from vaginal to placental samples during 16S rRNA gene sequence library generation.

Furthermore, *Lactobacillus* ASVs were also more prominent in samples of placental tissues of maternal origin, such as the decidua or basal plate, than placental tissues of fetal origin, such as the amnion, chorion, or villous tree. After separating placental sample data from non-labor term cesarean deliveries by fetal and maternal origin, with the exception of the Olomu et al. (39) study, *Lactobacillus* ASVs were absent from placental samples of fetal origin **(Table S2)**. In contrast, among samples of maternal origin from the Theis, Winters et al. dataset (86), *Lactobacillus* was the most relatively abundant ASV even after removal of likely DNA contaminants with DECONTAM, and in the Lauder et al. (37) study, only the maternal side of the single cesarean delivered placenta had a high predominance of *Lactobacillus*.

### The bacterial ASV profiles of placental samples and background technical controls cluster based on study origin

Beta diversity between placental samples and technical controls was visualized through Principal Coordinates Analysis for each study in the re-analysis to assess the extent of influence of background DNA contamination on the bacterial ASV profiles of placental samples (**Figure 3**). A majority of placental samples cluster with their respective technical controls across the studies. Specifically, in five of eleven studies, technical controls covered the entire bacterial profile spectrum of placental samples (**Figure 3A-E**), and in the remaining six studies which included technical controls, the bacterial profiles of most placental samples largely clustered with those of technical controls (**Figure 3F-K**). Placental samples in the latter group which did not cluster with technical controls were characterized by a predominance of *Lactobacillus* **(3F-H)**, *Cutibacterium* **(3I,K)**, *Gardnerella* **(3F)**, *Pseudomonas* **(3F)**, *Ureaplasma* **(3G)**, *Lactobacillus* **(3G-H)**, *Mesorhizobium* **(3I)**, *Prevotella* **(3J)**, *Actinomyces* **(3J)***, Streptococcus* **(3J)**, *Veillonella* **(3J)**, and *Staphylococcus* **(3K)**. Notably, the bacterial profiles of most placental samples from term cesarean deliveries were not significantly different from those of technical controls in either dispersion or structure **(Table S3)**. In cases where the structure of the bacterial profiles differed between placental samples and technical controls, but the dispersion of the bacterial profiles did not, it was only the bacterial profiles of the exterior surfaces of the placenta which differed from those of controls. In these cases, the bacterial profiles of the exterior surfaces of placental samples were characterized by *Cupriavidus, Serratia*, *Corynebacterium*, and *Staphylococcus* **(Table S3)**, the latter two of which are well-known commensal bacteria of the human skin (90).

**Figure 3.**
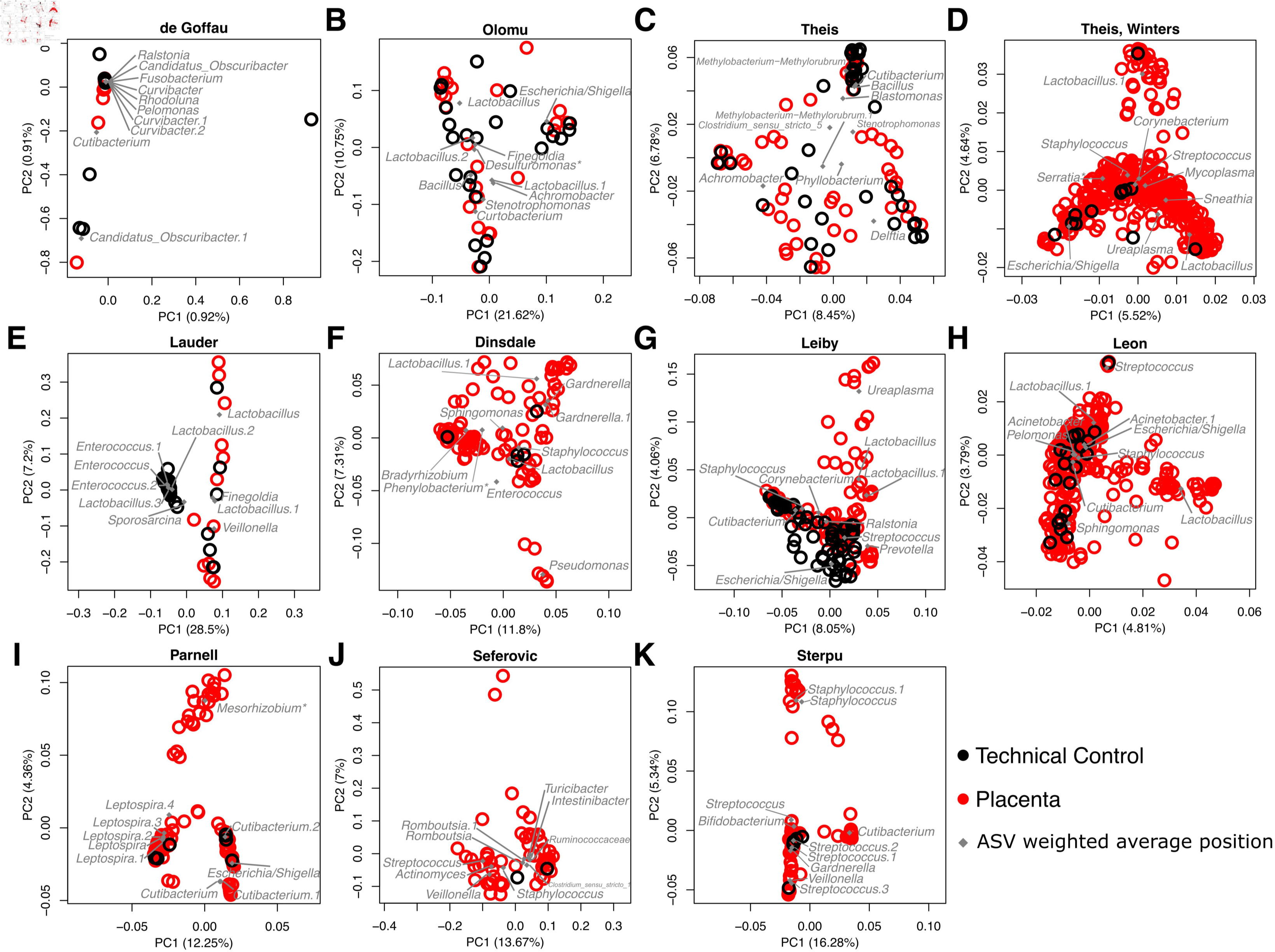
Principal Coordinates Analyses of the beta diversity of bacterial DNA profiles between placental samples and technical controls in published placental microbiota studies. Studies were included if technical controls were sequenced and made publicly available to account for background DNA contamination. Beta diversity between placental (red open circles) and technical control (black open circles) samples is illustrated by study in PCoA plots based on the Bray-Curtis dissimilarity index. Genus level classifications of the top ten ASVs in placental samples and technical controls by total reads are plotted at their weighted average positions (grey diamonds). Asterisks indicate ASV sequence genus level classifications which were assigned by NCBI BLAST with the highest percent identity in excess of 95%.

### The bacterial ASV profiles of vaginally delivered placental samples also cluster with their respective vaginal samples across studies

Six studies (29, 37–39, 50, 57) in the re-analysis included vaginal or vaginal-rectal swab samples as a complement to placental samples; four of these studies also included technical controls (37–39, 50). While most technical controls did not cluster with vaginal samples, placental samples typically clustered with vaginal samples and/or technical controls (**Figure 4A-D**), or if technical controls were not included in the study, with vaginal samples (**Figure 4E-F**). Notably, nine *Lactobacillus* ASVs were shared between the top ranked ASVs of placental and vaginal swab samples across five studies (37-39, 50, 57) (**Figure 4**).

**Figure 4.**
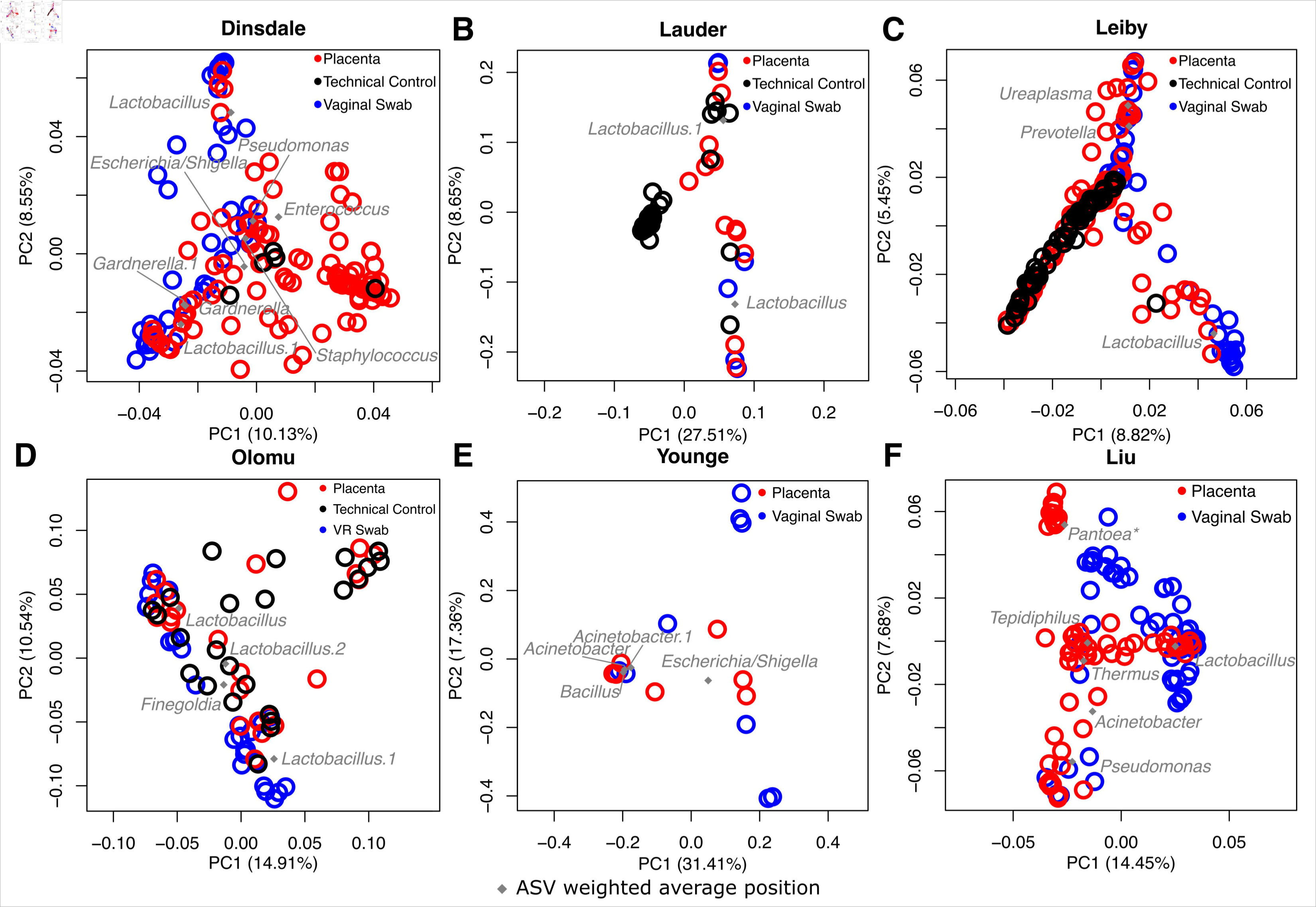
Principal Coordinates Analyses of the beta diversity of the bacterial DNA profiles of placental and vaginal/vaginal-rectal samples in placental microbiota studies. Prior published studies were included if vaginal or vaginal-rectal samples were sequenced and made publicly available alongside placental samples. The top ten ASVs shared between placental samples and technical controls, and the top ten ASVs in vaginal samples are plotted at their weighted average positions in the ordination space (grey diamonds) and their genus level classifications are noted. Agglomerated genus level classifications were plotted for the Liu dataset instead of ASVs since no ASVs were greater than 1% mean relative abundance across placental samples. Asterisks indicate ASV sequence genus level classifications which were assigned by NCBI BLAST with the highest percent identity in excess of 95%.

### Placental and technical control samples co-cluster by study and placental samples additionally cluster by mode of delivery

In order to fully utilize the capacity for ASVs to be directly compared across placental microbiota studies, taxonomy and ASV count tables were merged based on the exact ASV sequence data for six (29, 39, 41, 44, 50, 86) of seven studies (29, 39, 41, 43, 44, 86) which sequenced the V4 hypervariable region of the 16S rRNA gene using the PCR primers 515F and 805R. Principal Coordinates Analysis (PCoA) illustrated that placental and technical control samples formed distinct clusters based on study origin (**Figure 5A**; NPMANOVA using Bray-Curtis; placental samples: F=16.0, P=0.001; technical controls: F=4.64, P=0.001). The only exception was the Theis, Winters et al. dataset (86), which encompassed the bacterial profiles of placental samples from the other studies. This was likely due to the inclusion of samples in Theis, Winters et al. (86) from multiple regions of the placenta (i.e., amnion, amnion-chorion interface, subchorion, villous tree, and basal plate) as well as placentas from term and preterm vaginal and cesarean deliveries (**Figure 5A**). When stratifying by study and thereby taking study origin into account, placental and technical control samples did exhibit distinct bacterial DNA profiles (**Figure 5A**; F=6.66; P=0.512). When technical controls were excluded from the PCoA, discrete clustering of placental samples by study origin was still apparent (**Figure 5B**).

**Figure 5.**
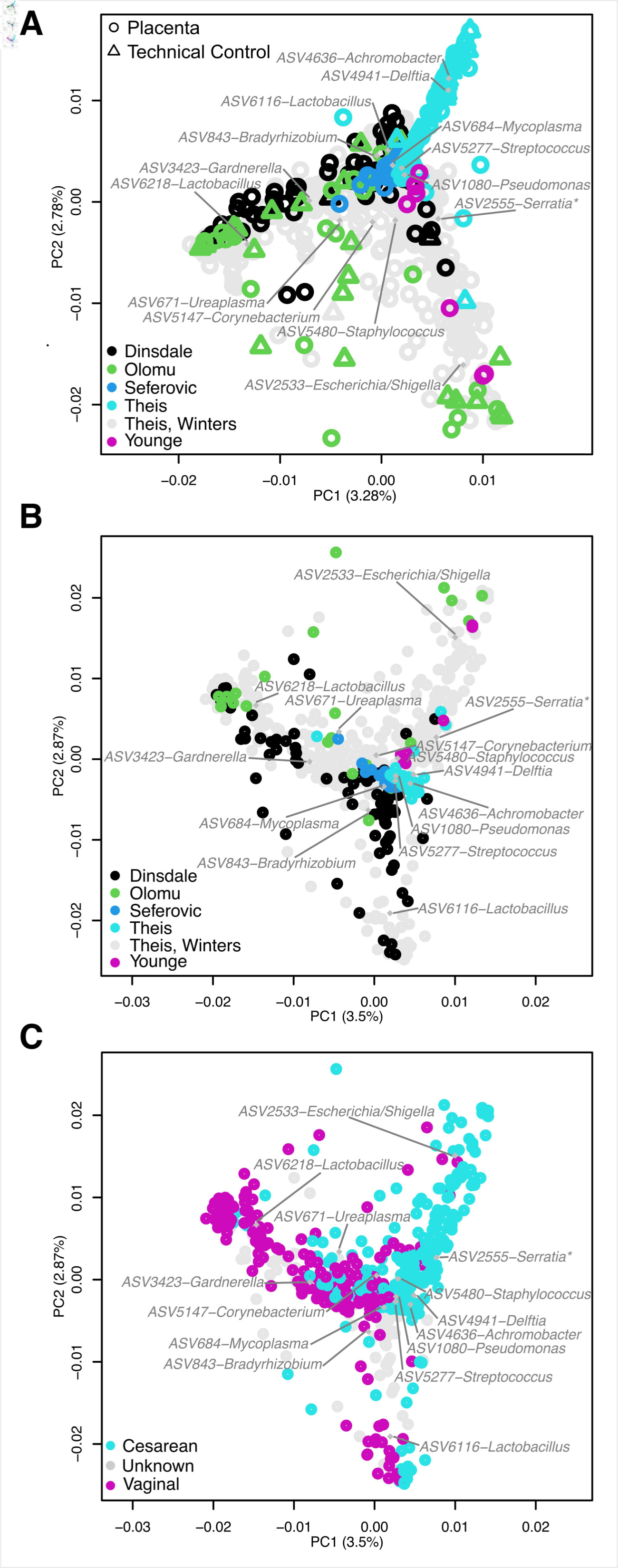
Placental and technical control samples cluster by study origin, mode of delivery, and gestational age at delivery. A) Beta diversity between placental (open circles) and technical control samples (open triangles) from studies which sequenced the V4 hypervariable region of the 16S rRNA gene is visualized through principal coordinate analysis (PCoA) based on the Bray-Curtis dissimilarity index. B) Beta diversity of placental samples without technical control samples from each study. C) Placental samples from the same six studies were characterized by mode of delivery and gestational age at delivery. Weighted average positions of ASVs greater than 1% were plotted as grey diamonds and labelled with genus level classifications. Asterisks indicate ASV sequence genus level classifications which were assigned by NCBI BLAST with the highest percent identity in excess of 95%.

Furthermore, the bacterial DNA profiles of placental samples were clearly affected by mode of delivery across studies (**Figure 5C**; F=21.6, P=0.001). Unsurprisingly, common vaginal bacteria such as *Lactobacillus*, *Ureaplasma*, and *Gardnerella* were predominant in the profiles of placental samples from vaginal deliveries (**Figure 5C**).

### Bacterial profiles of placental and technical control samples characterized using the V4 hypervariable region of the 16S rRNA gene share prominent ASVs

While placental samples from each study exhibited characteristic patterns of predominant ASVs, some ASVs such as ASV2533-*Escherichia/Shigella*, ASV6218-*Lactobacillus*, and ASV6216- *Lactobacillus* were predominant in the datasets of several studies **(Figure 6A-B, E)**. However, across studies, nearly every ASV that was consistently predominant in the bacterial DNA profiles of placental samples, was also consistently predominant in the profiles of the technical control samples from the same dataset (**Figure 6**). For instance, in two studies, all ASVs with a mean relative abundance greater than 1% in placental samples were also greater than 2% mean relative abundance in technical control samples (**Figure 6B-C**). In a third study, all ASVs other than ASV5229-*Cutibacterium* were also greater than 2% mean relative abundance across technical control samples (**Figure 6D**). These data collectively indicate that prominent placental ASVs were likely derived from background DNA contamination captured by the technical control samples.

**Figure 6.**
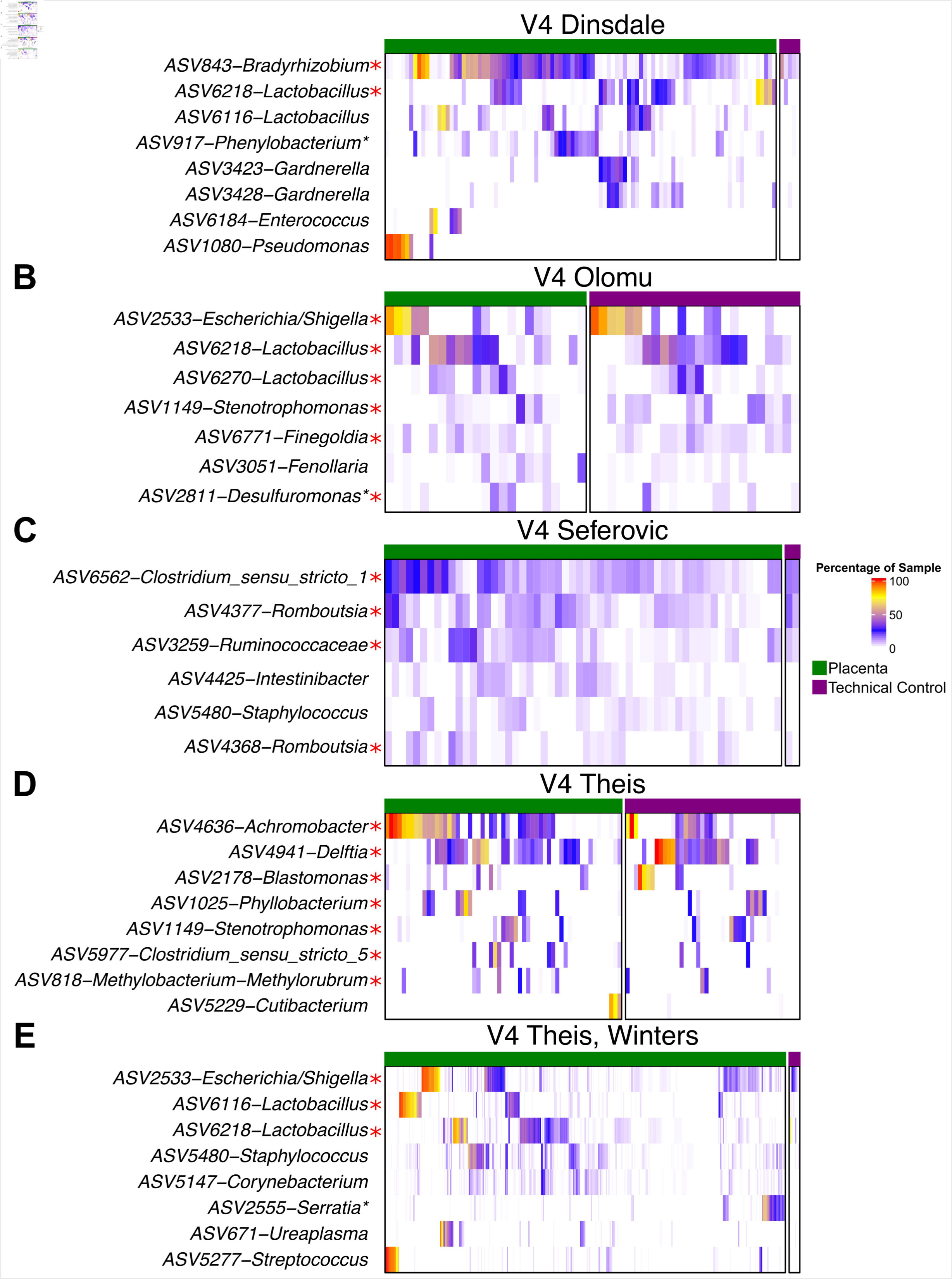
Heatmaps of the bacterial DNA profiles of placental and technical control samples from studies which sequenced the V4 hypervariable region of the 16S rRNA gene demonstrating a high degree of overlap between these two sample types. ASVs are listed by study if they had a mean relative abundance greater than 1% across placental samples (green bar). Red asterisks indicate ASVs which had a mean relative abundance greater than 2% across all technical control samples (purple bar) from that study. Regular asterisks indicate ASV sequence genus level classifications which were assigned by NCBI BLAST with the highest percent identity in excess of 95%.

## DISCUSSION

### Principal findings of the study

In this re-analysis of fifteen placental microbiota studies, of the ASVs which were ranked in the top five ASVs for relative abundance in any one study, *Lactobacillus* ASVs were clearly the most prevalent across studies. Yet, *Lactobacillus* ASV prevalence was explained by background DNA contamination, contamination from the birth canal during vaginal delivery, or well-to-well contamination from vaginal samples during the sequence library build process.

Overall, the bacterial DNA profiles of placental samples were highly similar to those of technical controls in their respective studies. Indeed, a secondary analysis of the six studies which targeted the V4 hypervariable region of the 16S rRNA gene for sequencing, showed that the bacterial DNA signal of both placental and technical control samples clustered by study of origin rather than by sample type. In addition, the top two ASVs in placental samples from each of the six studies in the secondary analysis were also the top ranked ASVs in technical controls from the corresponding study. Considered in isolation, placental samples clustered by mode of delivery, suggesting that the process of delivery greatly affected the bacterial DNA profiles of placentas.

Therefore, placental samples included in this re-analysis do not provide evidence of a consistent bacterial DNA signal in typical term pregnancy independent of mode of delivery. Instead, the modest consistency in bacterial DNA signals identified across studies was associated with general background DNA contamination or contamination introduced during vaginal delivery.

### The findings of this study in the context of prior reports

Currently, the extent of bacterial presence within the placenta is under debate. There have been numerous reviews, commentaries, and editorials, which have sought to synthesize and resolve conflicting results regarding the existence of a placental microbiota (3, 30, 32, 33, 66–68, 73, 91- 135). Although there has been disagreement about the existence of a placental microbiota in typical human pregnancy, there is a consensus that any given body site, including the placenta, can be at least transiently infected by microorganisms. Several reviews have emphasized that microorganisms in placental tissue would not be able to survive for long durations given the structure of the placenta and the immunobiological response of the host (3, 101). In contrast, some have proposed that microorganisms could survive intracellularly within the basal plate of the placenta and thus effectively evade the host immune system (68, 136). Many reviews addressing prior microbiota datasets have been challenged to draw conclusions given the multiple confounding factors which could significantly influence results: the specific 16S rRNA gene hypervariable region targeted for sequencing, brand and lot number of the DNA extraction kits, gestational age at delivery and sampling, mode of delivery of the placenta, inadequate metadata for deposited sequence data, and a general lack of technical controls to account for background DNA contamination. Regardless, many have viewed the current evidence for placental and/or *in utero* colonization as theoretically tenuous given the existence of germ-free mammals and the strong potential for background bacterial DNA to influence DNA sequencing surveys of low microbial biomass samples (36-41, 81, 103). Finally, similar to the results presented here in this re-analysis, the prevalence of *Lactobacillus* across placental samples in prior studies has been acknowledged, yet so too has been the high variability in the bacterial taxa reported within placental tissues across studies. Indeed, variability is high even across studies of similar cohorts from the same research groups (27, 41, 44, 63–65, 86). The current study sought to remedy the lack of consensus in the literature regarding the existence of a placental microbiota in typical term pregnancy through a re-analysis of the current publicly available data on placental microbiotas while controlling for targeted region of the 16S rRNA gene for sequencing, background DNA contamination, and mode of delivery.

### Mode of delivery must be taken into account when investigating the existence and structure of a placental bacterial DNA signal

Eleven studies (27, 38, 43, 44, 52, 54, 57, 59) concluded that the bacterial DNA signals in placentas from cesarean deliveries were not significantly different from those in placentas delivered vaginally. Yet, six other studies (36, 38, 42, 51, 86, 89) have reported that the bacterial DNA signals in placentas from vaginal and cesarean deliveries significantly differ. The latter studies have reported increased prevalence and relative abundance of *Lactobacillus* and other vaginally associated taxa in placentas from vaginal deliveries. Additionally, even the rupture of membranes, a prerequisite for labor and vaginal delivery, provides microorganisms access to the amniotic cavity (137) and thus the placenta, with prolonged access leading to microbial invasion and infection (138, 139). Notably, bacterial culture of placentas from vaginal deliveries have significantly higher positivity rates (18, 86), higher total colony counts (40), and a higher prevalence of bacterial colonies from *Lactobacillus* and *Gardnerella*, both of which are typical residents of the human vagina (78). In contrast, placentas from cesarean deliveries consistently yield bacteria which typically predominate on the skin, such as *Propionibacterium*, *Staphylococcus*, and *Streptococcus* (40, 90).

Importantly, through robust analysis of the entire bacterial DNA signal from hundreds of placental samples, this re-analysis clearly highlights the influence of mode of delivery on the bacterial DNA signal from placental samples by demonstrating mode of delivery-associated clustering across six studies. Furthermore, it is apparent that removing the exterior layers (i.e., amnion, chorion, and basal plate) of a placenta delivered vaginally is not sufficient to eliminate delivery associated DNA contamination of the sample since the diversity and structure of bacterial DNA profiles from the inner layers (i.e., subchorion, villous tree) of the placenta differed significantly between cesarean and vaginal deliveries. Evidence in the literature combined with this re-analysis warrants careful consideration of mode of delivery and even time since rupture of membranes (52, 138, 139) when investigating the bacterial DNA signal from placental samples.

### Background bacterial DNA limits analysis of bacterial 16S rRNA gene signal from the placenta

Theoretically, a low bacterial biomass community is detectable using 16S rRNA gene sequencing when its concentration is at least 10-100CFU/mL (140). However, discerning a true tissue-derived low bacterial DNA signal from other potential sources is exceedingly difficult.

This re-analysis, along with eight other studies (36-41, 81, 86), found that placental samples and technical controls share highly abundant bacterial taxa when 16S rRNA gene sequencing is used. Since technical controls represent the environment and reagents to which the placenta is exposed post-delivery, it follows that a majority of the bacterial DNA signal from placental samples is also acquired from those environments and reagents. While a placental tissue limit of bacterial detection through DNA sequencing is yet to be determined, other low-bacterial-biomass sample types such as oral rinse, bronchoalveolar lavage fluid, and exhaled breath condensate were predominated by stochastic noise below 10^4^ 16S rRNA gene copies per sample (141). Even the bacterial DNA signal from a pure culture of *Salmonella bongori* serially diluted to a final concentration of 10^3^ CFU/mL was mostly contamination (74). If these limits are comparable to those in placental tissue, then stochastic noise and background DNA contamination would predominate the bacterial DNA signal from placental tissue leaving any true DNA signal well beyond the detection limits of 16S rRNA gene sequencing. Therefore it follows that 16S rRNA gene sequencing is inadequate to make a clear assessment of the existence of a placental microbiota.

### Prior reports of 16S rRNA gene sequencing on placentas from term pregnancies

With the prior considerations in mind, out of the 40 studies which performed 16S rRNA gene sequencing on placental samples, 32 included at least some term deliveries. However, only 16 focused exclusively on placentas from term deliveries (28, 37, 39–41, 43, 49, 53, 54, 56–58, 62- 65). Additionally, only nine of these studies focused exclusively on placentas from cesarean deliveries (28, 39, 41, 49, 56, 58, 62, 64, 65) and only three included technical controls and their DNA sequencing data thus accounting for gestational age, mode of delivery, and background DNA contamination (39, 41, 49). Two of three concluded that there was no evidence for a placental microbiota in the context of term cesarean delivery (39, 41).

Many studies have reported evidence for a low biomass placental microbiota (27, 29, 30, 43–47, 49, 50, 52–54, 57, 58, 60, 61, 63–65, 82, 83, 136, 142) but only nine of these studies exclusively sampled placentas from term deliveries (43, 49, 53, 54, 57, 58, 63–65). Predominant bacterial taxa reported in these studies included *Pseudomonas* (54, 64, 65), *Lactobacillus* (49, 54), Bacteroidales (64), *Enterococcus* (63), *Mesorhizobium* (43), *Prevotella* (58), unclassified Proteobacteria (57), *Ralstonia* (43), and *Streptococcus* (54). Two studies from this term delivery subset, which sampled multiple regions of the placenta, observed gradients of *Lactobacillus* relative abundance across levels of the placenta, but in opposite directions (43, 49).

In contrast, five studies did not find evidence for a microbiota in placentas from term deliveries since neither the placental bacterial DNA signal from 16S rRNA gene sequencing (37, 39–41) nor the bacterial load as determined by quantitative real-time PCR (37, 39–41, 56) were significantly different from technical controls. One study even noted that no operational taxonomic units greater than 1% relative abundance in placental samples, were less than 1% in technical control samples, emphasizing the overlap between the two sample types (37). Three of these studies (40, 41, 56) also attempted to culture viable bacteria from placental tissue, but were rarely successful. In cases where culture was successful, viable bacteria often conflicted with the DNA sequencing results suggesting that cultured bacteria were likely contaminants (40, 41).

### Clinical significance

#### Non-viable or viable bacterial DNA could feasibly be filtered from maternal blood by the placenta leading to a placental bacterial DNA signal

The placenta is a transient internal organ with functions that include promotion of gas exchange, nutrient and waste transport, maternal immunoglobulin transport, and secretion of hormones critical for fetal growth and development (143). These exchanges and transfers occur due to diffusion gradients between fetal and maternal blood, the latter of which bathes the chorionic villi in the intervillous space of the placenta (99). This maternal blood, which cannot be fully drained from the placenta before biopsy or sampling, can undoubtedly contain bacterial particles or even the remnants of a low-grade bacterial infection (56, 103, 144). Because of its structure, the placenta functions as a filter and retains these particles or bacteria, dead or alive. A bacterial DNA signal due to this filtering process would be extremely weak and transient. In addition, the bacterial taxa identified would be highly variable since they do not correspond to a specific niche, which is consistent with the findings of this re-analysis.

#### Infection is a potential source for the placental bacterial DNA signal

Instead of *in-utero* colonization, it is more likely that the bacterial DNA signal coming from a subset of placental samples is caused by infection. It is curious to note that specific bacteria are associated with stronger bacterial DNA signals and inflammation in placental tissue resulting in adverse pregnancy outcomes including preterm birth and/or preterm prelabor rupture of membranes (PPROM) (52, 55, 89). Spontaneous preterm birth has been shown to increase bacterial load (55) and the relative abundances of several taxa in placental samples including but not limited to *Ureaplasma* (26, 36, 38, 42, 51, 52, 145, 146), *Fusobacterium* (51, 52), *Mycoplasma* (42, 51, 52), *Streptococcus* (36, 51), *Burkholderia* (27), *Escherichia/Shigella* (55), *Gardnerella* (51), *Gemella* (52), and *Pseudomonas* (50). *Ureaplasma urealyticum*, *Mycoplasma hominis*, *Bacteroides* spp., *Gardenerella* spp, *Mobiluncus* spp., various enterococci, and *Streptococcus agalactiae* (also known as Group B Streptococcus or GBS) are frequently associated with histologic acute chorioamnionitis as well as uterine infection (16, 26, 99, 146). GBS is also a major cause of early onset neonatal sepsis and has been commonly isolated at autopsy in addition to *E. coli*, and *Enterococcus* (16, 147). While strain level variation could conclusively determine the pathogenicity of bacterial DNA in the placenta, 16S rRNA gene sequencing is not capable of such resolution. Nevertheless, the DNA of the notoriously pathogenic bacterial genera detailed above were all found in placental tissue, suggesting an invasive phenotype rather than commensal colonization.

### Recommendations for future studies

In order to establish the existence of a viable placental microbiota several criteria need to be met, which have been detailed previously (36, 41). Studies which aim to assess the viability of a bacterial DNA signal in a purported low biomass sample type should start with the null hypothesis that the entire DNA signal results from contamination and subsequently attempt to reject it with experimental evidence (148). Therefore, any study evaluating a potential microbiota of the placenta should attempt to demonstrate viability through both culture and DNA sequencing. Placentas should come from term cesarean deliveries without labor to obviate contamination during vaginal delivery and subjects should be screened to ensure that only healthy women are sampled (i.e., no history of antenatal infection, pre-eclampsia, recent antibiotic use, signs of infection or inflammation). Additionally, future studies should include ample sequenced technical controls in order to identify and account for sources of contamination, which will inevitably exist no matter how rigorous and/or sterile the protocol (71). Further, biological replicates from the same placenta should also be included to ascertain the consistency of any bacterial DNA signal. Since 16S rRNA gene sequencing limits of detection have not yet been thoroughly interrogated in placental tissue, serial dilutions of spiked-in live bacteria or cell-free DNA should be included in a portion of tissue samples to demonstrate the feasibility of recovering the bacterial DNA signal from placental tissue. When multiplexing samples, unique dual index primer sets should be used to eliminate the possibility of barcode hopping which is another source of sample “contamination” (149, 150), and before sequencing, low biomass samples should be segregated from higher biomass samples to avoid well-to-well contamination (39, 151). Finally, in conjunction with publishing, all sequence data and detailed metadata should be submitted to a public database so that others can replicate the work and verify the results.

### Strengths of this study

Broad searches of the available literature were utilized to ensure that all publicly available 16S rRNA gene sequencing data of placental samples (with associated metadata to partition pooled sample data) were incorporated into the re-analysis, which re-examined the data with in-depth comparisons of term placental samples to technical controls. This allowed for the detection of background DNA contamination in the bacterial DNA signal from placental tissue. In addition, potential confounding variables such as mode of delivery, gestational age at delivery, and 16S rRNA gene target hypervariable region were controlled for whenever possible. By utilizing DADA2 to process the sequence data, variation and biases due to post-sequencing processing were eliminated. This enabled ASV-to-ASV comparisons for six studies which targeted the same 16S rRNA gene hypervariable region using the same PCR primers, a first in the placental microbiota field.

### Limitations of this study

The quality and public availability of data and metadata were the primary limiting factors of this re-analysis. Unfortunately, the availability of metadata or even the data themselves is a pervasive issue in the microbiome field (152–154). While study cohort statistics were well reported overall, detailed metadata for each subject are required in order to perform a proper re-analysis. Ideally, any study investigating the existence of a viable placental microbiota would, at a minimum, include associated metadata by subject for potential confounders (e.g., gestational age at delivery, and mode of delivery).

An additional limitation of the current study was that the impacts of individual low abundance ASVs (i.e., less than 1% mean relative abundance) were not evaluated, but these ASVs were likely stochastic environmental DNA contamination. Finally, while the R package DECONTAM was used to remove likely contaminants by comparing the prevalence of ASVs in biological samples and technical controls, this tool is not appropriate for identifying contaminants introduced during sampling or delivery. In addition, the contaminant identification accuracy of DECONTAM also diminishes when used on low biomass samples such as placental samples where the majority of the sequences are likely contaminants (71, 155).

## Conclusion

The initial premise of this critical review and re-analysis was to determine if a true consistent bacterial DNA signal could be identified in placental samples from women delivering at term across studies despite various differences in sampling methodologies and sequencing analyses. 16S rRNA gene sequencing data from fifteen studies were processed and analysed in the same manner to control for as much post-sequencing variation as possible. By doing so, *Lactobacillus* ASVs were identified as the most prevalent top-ranked ASVs by relative abundance across studies; however, their prevalence in placentas from term cesarean deliveries was attributable to some form of contamination in every case. While bacterial DNA signals were observed in placental samples, they were largely similar to those seen in technical control samples.

Furthermore, the bacterial DNA signal from placental samples clustered by mode of delivery, indicating placental delivery-associated contamination. This observation, in combination with the existence of germ-free mammals (156, 157), has yet to be reconciled with *in-utero* colonization. Even if the placenta has a bacterial DNA signal apart from that of background DNA or delivery-associated contamination, alternative sources for the bacterial DNA signal such as extracellular DNA or dead bacteria circulating within maternal blood still need to be ruled out.

As we push the boundaries of DNA sequencing technologies we need to tread carefully, especially in purported low-biomass sites such as the placenta. The limitations of current DNA sequencing technology make detection of a legitimate signal or determination of viability unattainable at such low levels (72, 74). Regardless, a bacterial DNA signal can indeed be detected even in placentas from term cesarean deliveries, but the placenta is unlikely to be the source. Only after demonstrating a valid, viable bacterial DNA signal from term cesarean deliveries, through sterile protocol, with technical controls, and associated culture positive data, can we evaluate the degree to which the maternal immune system tolerates these bacteria and whether their presence resembles commensal existence or infection. Finally, the placental microbiota may or may not exist, but it is quite clear that attempts to maintain sterility and avoid contamination have not been successful since the vast majority of sequencing reads from placental samples can be attributed to multiple modes of contamination. Therefore, sequencing methodologies require significant improvement before a placental microbiota can be established as 16S rRNA gene sequencing appears to lack the ability to discriminate between a markedly low biomass microbiota and background DNA contamination at present.

## MATERIALS AND METHODS

### Study inclusion criteria

Searches for “human placental microbiome”, “placenta microbiota”, “placental microbiomes”, and “placenta 16S” were queried on PubMed with a cutoff date of 6/16/21 to identify studies addressing the existence of a placental microbiota or lack thereof. Additionally we included our recent preprint (86) in this pool of studies. Of the 387 unique studies identified, 58 performed primary research and 41 implemented 16S rRNA gene sequencing on placental samples (Error! Reference source not found.). Therefore we focused on 16S rRNA gene sequencing data. 16S rRNA gene sequencing is a well-characterized way of detecting and classifying bacterial communities within biological samples (158–160), and it is potentially sensitive enough to detect the typically low number of 16S rRNA bacterial gene copies hypothesized to be in the placenta (34, 161). 33 of the 41 studies which implemented 16S rRNA gene sequencing included at least one placental sample from an uncomplicated delivery at term (27-29, 36-45, 47, 49-65, 86, 142). However, only 15 of these 33 studies included publicly available 16S rRNA gene sequence data (i.e., sequencing files uploaded to a public database with a published and accurate accession number with sufficient metadata to partition pooled sample data) (29, 36–44, 50, 53, 57, 58, 86). Thus, the re-analysis ultimately included 15 studies.

### Processing of 16S rRNA gene sequence data using DADA2

Fastq files of the 16S rRNA gene sequence data from samples included in each study were downloaded from publicly accessible databases. If a study included fastq files that contained sequence data from multiple samples, the data were demultiplexed using QIIME2 (version 2020.2) (162) and SED (GNU Sed 4.7), a stream editor for text processing (163).

Sequence datasets from each study were individually processed using the Differential Abundance Denoising Algorithm (DADA2), which is an R package designed to partition 16S rRNA gene sequences into distinct Amplicon Sequence Variants (ASVs) and to taxonomically classify the resultant ASVs (87). R version 3.6.1 (164) was used for DADA2 processing and all downstream analyses. Processing followed the 1.16 DADA2 guidelines (https://benjjneb.github.io/dada2/tutorial.html), except when stated otherwise. Samples that had an average sequence quality score which dipped below 30 before the expected trim length cutoffs were removed from the dataset. Trim length cutoffs were set to maximize the read length and number of passing samples while still removing poor quality base calls from the ends of reads.

Reads were then filtered using the filterAndTrim() function with *multithread* set to TRUE to enable parallel computation and decrease real time spent computing. Error rates of base calling in the filtered sequences were inferred from the data using the learnErrors() function with *multithread* set to TRUE. Using the inferred error rates, sequences were partitioned into ASVs with *pool* and *multithread* set to TRUE. If the dada() function failed to complete partitioning after 7 days for a particular study’s dataset, which occurred for only one study (36), *pool* was set to FALSE for sample independent sequence partitioning.

If forward and reverse sequences were not yet merged, they were merged using the mergePairs() function. In cases where the forward and reverse reads were already merged in publicly available data files, the DADA2 merging step was omitted and the code adjusted for merged input sequences. Merged sequences with lengths greater or less than 20 nucleotides from the expected amplified region were removed from the data set since they were most likely due to non-specific merging of forward and reverse reads resulting in extra-long or extra-short reads.

Chimeric sequences were detected and removed using the removeBimeraDenovo() function with *multithread* set to FALSE. This employs the default consensus method instead of the pooled method. The consensus method determines chimeric sequences in each sample and then compares detected chimeric sequences across samples for a consensus. Taxonomy was assigned to sequences using the Silva 16S rRNA gene bacterial database (v 138) (159, 165). Species assignments were added, when possible.

For each study, merged datasets of ASV counts and taxonomic classifications were filtered using functions from the R package dplyr (166) to remove ASVs that were classified as mitochondrial, chloroplast, or not of bacterial origin. ASVs not classified at the phylum level and samples which did not have at least 100 sequence reads after filtering were removed from the data set.

### Removal of likely DNA contaminants through the R package DECONTAM

To control for background DNA contamination, the R package DECONTAM was used to identify and remove sequences which were more prevalent in technical controls than in placental samples. For likely sequence contaminant removal, studies which included at least six technical controls (36-43, 50, 86) were included based on the recommendation of the authors of DECONTAM (88). Technical controls included air swabs, blank DNA extraction kits, and template-free PCR reactions. The DECONTAM function isNotContaminant() was used to remove ASVs which were more prevalent in technical controls than in biological samples.

Thresholds were study specific with the goal of excluding most of the low prevalence ASVs while retaining high prevalence ASVs not likely to be contaminants. Despite using these stringent study specific thresholds, the results were unchanged if the default threshold of 0.5 was used instead.

### Normalization of 16S rRNA gene sequence datasets within and across studies

All datasets were normalized using the function rarefy_even_depth() from the R package phyloseq (1.30.0) (167). Following the normalization process, samples whose sequence libraries did not have at least 100 reads were excluded. The remaining samples were subsampled without replacement (i.e., the same sequence was never reselected when subsampling) to the minimum number of sequences per sample within a study. *RNGseed* was set to 1 to fix the seed for reproducible random number generation. This normalization approach was utilized since 16S rRNA gene read counts can vary by five orders of magnitude among samples in a single study. Given this degree of variability, normalization to the same sequence depth is justified and required for accurate comparisons of 16S rRNA gene profiles among samples (168).

### Data Visualization

Heatmaps illustrating the relative abundances of ASVs were prepared using the ComplexHeatmap R package (version 2.2.0) (169). Samples were grouped by sample type and ASVs were ordered based on ASV mean relative abundances within samples.

The function vegdist() from the R package vegan (version 2.5-6) (170) was used to create Bray-Curtis dissimilarity matrices which were then used as the basis for Principal Coordinates Analysis (PCoA) plots that were generated using the pco() function from the R package ecodist (version 2.0.7) (171). The Bray-Curtis index was used because it takes into account both the composition and structure of 16S rRNA gene sequence bacterial profiles (172). The Lingoes method was used to correct for negative eigenvalues so that dissimilarity between samples could be completely explained in Euclidean space (173).

All code to produce the published figures from the raw data is included in the supplementary materials in an R markdown file available at https://github.com/jp589/Placental_Microbiota_Reanalysis.

### Statistical analysis

Homogeneity of 16S rRNA gene sequence profiles between placental samples and technical controls was tested using betadisper() from the R package vegan (version 2.5-6) (170).

Differences in 16S rRNA gene profile structure between placental samples by sampling level and technical controls were evaluated using the function pairwise.adonis() from the R package pairwiseAdonis (version 0.4) (174).

All code to recapitulate these analyses are included in an R markdown file available at https://github.com/jp589/Placental_Microbiota_Reanalysis.

## ACKNOWLEDGEMENTS

Some raw data files and metadata included in the analyses were not publicly available, yet were generously provided upon request. Specifically, Drs. Shuppe-Koistinen and Hugerth provided raw sequence data for technical control samples for the Sterpu et al. study, as well as mode of delivery metadata, and Dr. Leon provided metadata for gestational age at delivery and mode of delivery for the Leon et al. study, which used samples taken from the Baby Bio Bank (175). We thank David Kracht and Dylan Millikin, both members of the Theis laboratory, for feedback and editing on the manuscript and the R markdown file.

## DATA DEPOSITION

Raw data for each of the studies included in analyses can be accessed through the following accession numbers:

De Goffau et al. PMID: 31367035 /Accession no. ERP109246;

Dinsdale et al. PMID: 33194782 /Accession no. PRJEB39698;

Gomez-Arango et al. PMID: 28240736 /Accession no. PRJNA357524;

Lauder et al. PMID: 27338728 / Accession no. PRJNA309332;

Leiby et al. PMID: 30376898 / Accession no. PRJNA451186;

Leon et al. PMID: 29776928 / Accession no. PRJEB25986;

Liu et al. PMID: 31685443 / Accession no. PRJNA559967;

Olomu et al. PMID: 32527226 / Accession no. PRJNA577959;

Parnell et al. PMID: 28894161 / Accession no. PRJNA395716;

Seferovic et al. PMID: 31055031 / Accession no. PRJNA511648;

Sterpu et al. PMID: 32871131 / Accession no. PRJEB38528;

Tang et al. PMID: 33193081 / Accession no. PRJNA564455

Theis et al. PMID: 30832984 / Accession no. PRJNA397876;

Theis, Winters et al. bioRxiv MS: 497119 / Accession no. PRJNA692425;

Younge et al. PMID: 31479427 / Accession no. PRJNA557826

## DATA AVAILABILITY STATEMENT

All DADA2 processed sequence data and metadata from the studies included in this critical review, as well as an R markdown file with the code to produce each of the figures and tables, are available online at https://github.com/jp589/Placental_Microbiota_Reanalysis. In addition, an R package ‘dada2tools’ with functions for efficient analysis of the data, is available at https://github.com/jp589/dada2tools.

